# Low- and high-level information analyses of transcriptome connecting endometrial-decidua-placental origin of preeclampsia subtypes: A preliminary study

**DOI:** 10.1101/2023.10.12.562143

**Authors:** Herdiantri Sufriyana, Yu-Wei Wu, Emily Chia-Yu Su

## Abstract

**Background:** Existing proposed pathogenesis for preeclampsia (PE) was only applied for early onset subtype and did not consider pre-pregnancy and competing risks. We aimed to decipher PE subtypes by identifying related transcriptome that represents endometrial maturation and histologic chorioamnionitis.

**Methods:** We utilized eight arrays of mRNA expression for discovery (*n*=289), and other eight arrays for validation (*n*=352). Differentially expressed genes (DEGs) were overlapped between those of: (1) healthy samples from endometrium, decidua, and placenta, and placenta samples under histologic chorioamnionitis; and (2) placenta samples for each of the subtypes. They were all possible combinations based on four axes: (1) pregnancy-induced hypertension; (2) placental dysfunction-related diseases (e.g., fetal growth restriction [FGR]); (3) onset; and (4) severity.

**Results:** The DEGs of endometrium at late-secretory phase, but none of decidua, significantly overlapped with those of any subtypes with: (1) early onset (*p*-values ≤0.008); (2) severe hypertension and proteinuria (*p*-values ≤0.042); or (3) chronic hypertension and/or severe PE with FGR (*p*-values ≤0.042). Although sharing the same subtypes whose DEGs with which significantly overlap, the gene regulation was mostly counter-expressed in placenta under chorioamnionitis (*n*=13/18, 72.22%; odds ratio [OR] upper bounds ≤0.21) but co-expressed in late-secretory endometrium (*n*=3/9, 66.67%; OR lower bounds ≥1.17). Neither the placental DEGs at first-nor second-trimester under normotensive pregnancy significantly overlapped with those under late-onset, severe PE without FGR.

**Conclusions:** We identified the transcriptome of endometrial maturation in placental dysfunction that distinguished early- and late-onset PE, and indicated chorioamnionitis as a PE competing risk. This study implied a feasibility to develop and validate the pathogenesis models that include pre-pregnancy and competing risks to decide if it is needed to collect prospective data for PE starting from pre-pregnancy including chorioamnionitis information.

## 1. Introduction

Preeclampsia (PE) is one of pregnancy-induced hypertension (PIH) subtypes related to placenta and endothelial dysfunction [1, 2]. This disease makes the survival more susceptible to cardiovascular diseases later in life [3]. Many studies have proposed pathogeneses for PE [4]. Most of these were typical for the early-onset subtype and shared with those of fetal growth restriction (FGR) [5], whereas both PE and FGR were placenta dysfunction-related diseases (PDDs). Meanwhile, the early-onset subtype only contributed to <30% cases of PE [6]. Therefore, regardless of numerous proposed pathogeneses, the common etiology for most of the PE subtypes is still unclear.

Worldwide, PE affected 3–8% pregnant women [7] and contributed to 11–18% maternal deaths. The risk of hypertension later in life increased 3.7 times for women with a history of PE and the onset was 7.7 years earlier with that of PIH, compared to women without PE or PIH, respectively [8]. Hypertension contributed to all-cause morbidities and mortalities in one fourth adults worldwide, although it is a modifiable risk factor [9]. This disease was more common in postmenopausal women compared to either men or the premenopausal counterparts and only 50% have controlled their blood pressure despite the well-awareness of necessary medications [10]. In addition, since the only cure is early delivery, PE was also the major contributor of prematurity and low-birth-weight infants [11], which led to neonatal deaths [12]. The preterm infants increased neonatal intensive care unit utilization and it was not reduced by the infants born from preeclamptic mother given preventive intervention using low-dose aspirin at 11–13 weeks’ gestation. Infants born from the preeclamptic mother also demonstrated signs of cardiac injury [13], which may increase risk of cardiovascular diseases later in life. Therefore, PE prevention has several impacts to mother and child healthcare, including the mortalities, morbidities, resources utilization, and cardiovascular diseases later in life.

Improvements of prevention strategy for any subtypes of PE need understanding of the pathogeneses. These were commonly believed to occur in the first trimester of pregnancy based on timing for the most successful prediction [14]. Yet, most of the comparisons were made against those conducted at the next trimesters without considering pre-pregnancy period [15]. Enormous theories have been proposed for the first-trimester pathogenesis, culminated into 2-stage theory [16, 17]. This consisted of two sequential dysfunctions in placenta and endothelium. However, the cause of pathophysiological derangement in placenta was still unclear [5]. Antecedents of this event were revealed by association between PE with either endometrial maturation [18] or metagenomics profiling of placenta [19]. There was a significant number of differentially expressed genes (DEGs) overlapped between those from preeclamptic chorionic villi sampling (CVS) and those from pathological endometrium [20]. But, the overlapped DEGs were regulated in the same direction instead of opposite ones, indicating the likelihood of co-occurrence instead of potential causal-effect relationship. Meanwhile, many publications describing association between microbiome and PE were proof-of-concept reviews instead of research articles. Eventually, PE remains a vascular disease with unknown etiology. This study aimed to identify transcriptome representing endometrial maturation and histologic chorioamnionitis, enriched by DEGs of the PE subtypes, using microarray meta-analysis workflow at low- and high-level information.

## 2. Materials and Methods

### 2.1. Dataset integration

A previous workflow on microarray dataset integration was applied [21]. We utilized 15 publicly-accessed microarray experiments of mRNA expression (*n*=653). The datasets were queried in Gene Expression Omnibus (GEO) and Array Express databases. To understand how these datasets helped in achieving the objective of our study, it is important to describe the spatial and temporal contexts of datasets in this study and the conditions they represented (Figure 1). Our datasets covered from pre-to post-pregnancy (post-partum) period. Pre-pregnancy period was represented by endometrial samples, while the pregnancy period was represented by either decidual (maternal side) or placental (fetal side) samples. The placental samples also represented the post-partum period in term of chronic/gestational hypertension phenotype, as defined by the original study. Gestational hypertension starts from 20 weeks’ gestation to 6 weeks after delivery, but this condition is considered as chronic hypertension if the elevated blood pressure persisted more than 6 weeks after delivery. Furthermore, our datasets also covered histologic chorioamnionitis, the PE subtypes, and hemolysis, elevated liver enzymes, and low platelets (HELLP) syndrome (mostly preceded PE, but may occur without PE).

**Figure 1.**
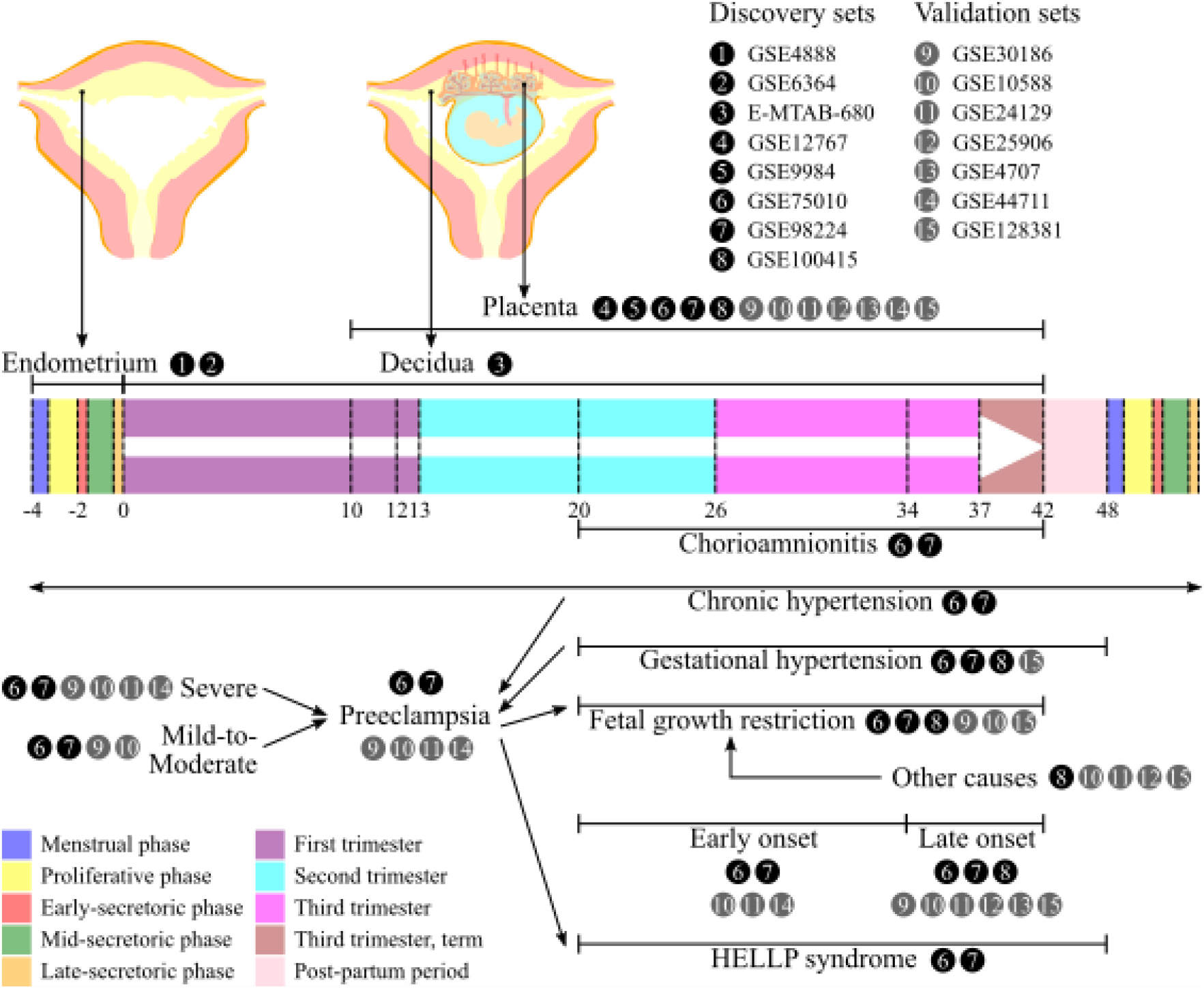
The spatial and temporal contexts of datasets in this study and the conditions they represented. HELLP, hemolysis, elevated liver enzymes, and low platelets.

We utilized the datasets 1 to 8 for DEGs discovery sets (Figure 1; Tables S1 and S2), which were GSE4888 (*n*=27; dataset 1) and GSE6364 (*n*=37; dataset 2) for endometrium, E-MTAB-680 (*n*=24; dataset 3) for decidua, GSE12767 (*n*=12; dataset 4) and GSE9984 (*n*=12; dataset 5) for CVS (i.e., the first-trimester placenta), and GSE75010 (*n*=157; dataset 6), GSE98224 (*n*=48; dataset 7), and GSE100415 (*n*=20; dataset 8) for third-trimester placenta of normotensive pregnant women and those with several subtypes of PE, other PIH, and other PDDs. The placental samples also consisted of those with and without either histologic chorioamnionitis or HELLP syndrome, but we included second-trimester placenta in addition to third-trimester placenta. All the discovery sets applied total RNA extraction. Endometrium datasets covered proliferative, and early-, mid-, and late-secretory phases. These four phases respectively represented endometrial maturation. Meanwhile, decidua datasets consisted of ectopic pregnancy (implantation site outside endometrium) without or with intermediate decidualization, and intrauterine pregnancy (implantation site inside endometrium) with intermediate and confluent decidualization. These four conditions represented decidualization from the lowest to the highest degree. For endometrium datasets, we excluded subjects with endometriosis and ambiguous histology reading of endometrial phases. For placenta datasets, we only included subjects with phenotypes that fitted the group definitions (see 2.2 Group definition). There were overlapped samples between GSE75010 and GSE98224 (*n*=48), but the duplicates were removed. There were no additional eligibility criteria applied for decidua and CVS datasets beyond those from the original datasets.

For validation sets (Figure 1; Table S1 and S2), we utilized the datasets 9 to 15, which were GSE30186 (*n*=12; dataset 9) with GPL10558 Illumina HumanHT-12 V4.0 expression beadchip, GSE10588 (*n*=43; dataset 10) with GPL2986 ABI Human Genome Survey Microarray Version 2, GSE24129 (*n*=24; dataset 11) with GPL6244 Affymetrix Human Gene 1.0 ST Array (transcript [gene] version), GSE25906 (*n*=60; dataset 12) with GPL6102 Illumina human-6 v2.0 expression beadchip, GSE4707 (*n*=14; dataset 13) with GPL1708 Agilent-012391 Whole Human Genome Oligo Microarray G4112A (Feature Number version), GSE44711 (*n*=16; dataset 14) with GPL10558 Illumina HumanHT-12 V4.0 expression beadchip, and GSE128381 (*n*=183; dataset 15) with GPL17077 Agilent-039494 SurePrint G3 Human GE v2 8x60K Microarray 039381 (Probe Name version). All the validation sets also applied total RNA extraction. Since most of the platforms were different, several DEGs might not be included in both discovery and validation sets corresponding the same subtype. Thus, we only used the intersected genes among them.

We did not apply data integration for discovery sets. The experiments using third-trimester placenta were conducted by the same microarray platform of GPL6244 Affymetrix Human Gene 1.0 ST Array (transcript [gene] version). Meanwhile, the remaining experiments using endometrium, decidua, and the first-trimester placenta were conducted by the other platform, which was GPL570 Affymetrix Human Genome U133 Plus 2.0 Array. Therefore, we identified DEGs separately for each tissue that used the same platform (Figure 2A).

**Figure 2.**
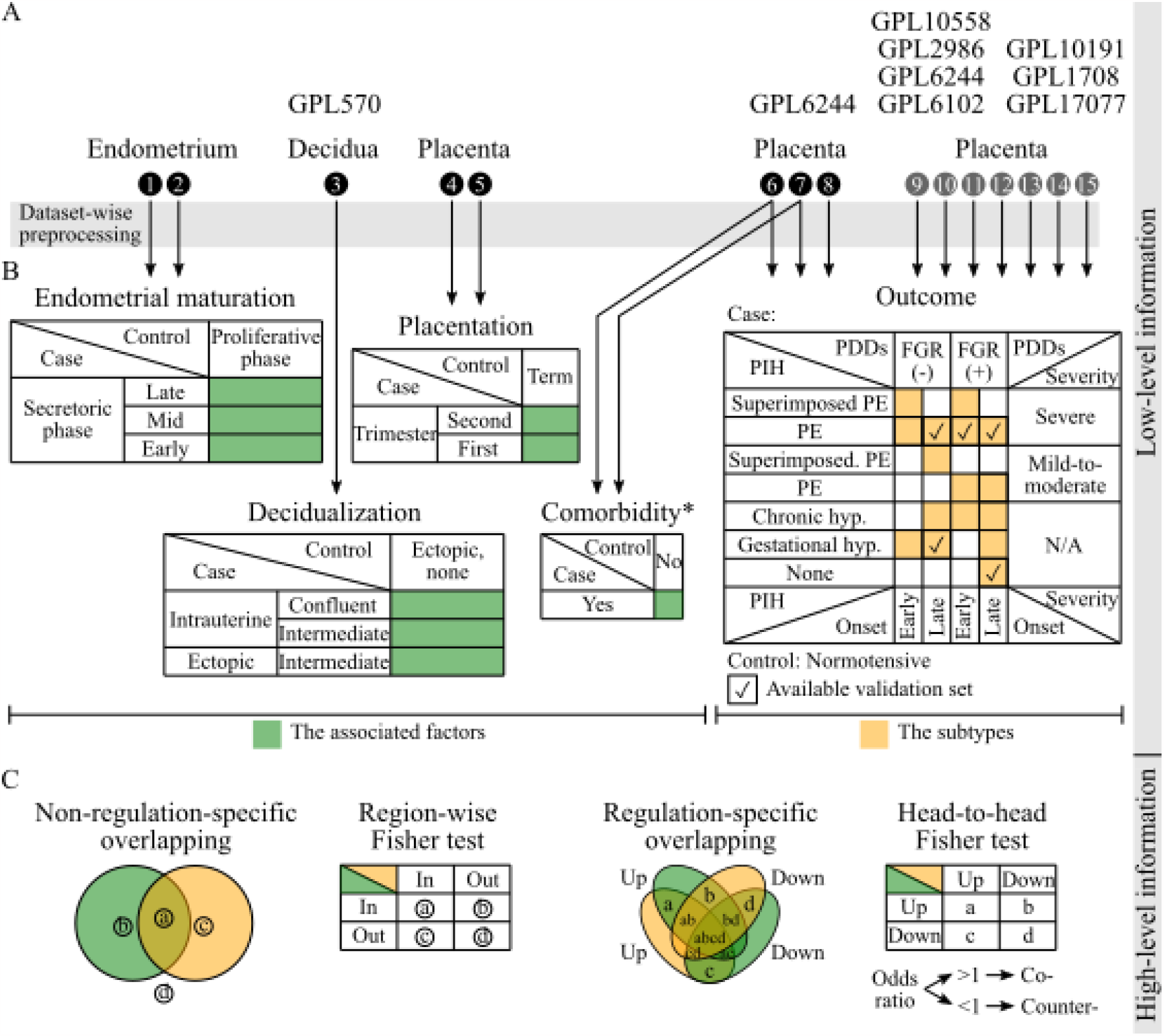
The analytical pipeline: (A) Data preprocessing, including quality control, for each dataset; (B) Differential expression analysis for discovery sets and data integration for validation sets; (C) Gene-set overlap analysis. *, conducted for chorioamnionitis/HELLP in either second or third trimester; FGR, fetal growth restriction; HELLP, hemolysis, elevated liver enzymes, and low platelets; hyp.; hypertension; N/A, not applicable; PDDs, placenta dysfunction-related diseases; PE, preeclampsia; PIH, pregnancy-induced hypertension.

Several pathogenesis models for PE subtypes will be developed in the extension work of this preliminary study according to the discovery sets without merging experiments by different platforms. To get comparable gene expression between discovery and validation sets after determining the DEGs and before developing the models, we normalized the validation sets according to the quantile distribution of the discovery controls, as previously described [22]. Therefore, the expression values are centered to those of the discovery controls. To ensure the comparable expression is achieved, we conducted principal component analysis. The samples were not separated among the experiment groups (Figure S1); thus, we could use the validation sets.

For the downstream analysis, the transcripts were summarized into genes. The raw expression data were combined according to the group definition. Outliers were estimated according to relative log expression before normalization and hierarchical clustering of sample-to-sample distances after normalization [23]. After removing outliers, the raw expression data were background-corrected and normalized using robust multi-array average algorithm. Quality control was conducted by data visualization using boxplot, quantile-to-quantile plot, and the MA plot, and confounder identification by surrogate variable analysis.

### 2.2. Group definition

PE is a syndrome characterized by both chronic/gestational hypertension and gestational proteinuria. PE subtypes in these datasets were all possible combinations based on four axes (Figures 1 and 2B): (1) PIH; (2) PDDs; (3) onset; and (4) severity. By PIH, there were two PE subtypes: (1) PE (i.e., gestational hypertension with gestational proteinuria); and (2) superimposed PE (i.e., chronic hypertension with gestational proteinuria). By PDDs, there were PE subtypes without and with FGR. By onset, there were two PE subtypes: (1) early onset (<34 weeks’ gestation); and (2) late onset (≥34 weeks’ gestation). By severity, there were two PE subtypes: (1) mild-to-moderate (i.e., systolic and diastolic blood pressures [SBP/DBP] of 140/90 to <160/<110 mm Hg with proteinuria of 300 to 2000 mg/24 h, and HELLP negative); and (2) severe (i.e., SBP/DBP of ≥160/110 mm Hg with proteinuria of >2000 mg/24 h, or HELLP positive). As comparators, we also included either chronic/gestational hypertension without gestational proteinuria, of which subtypes were also defined by PDDs and onset but not severity axes.

Using the distribution variances of gene expressions from controls in each tissue dataset, power analysis was conducted to estimate sample size for differential expression analysis with multiple testing by Benjamin-Hochberg false discovery rate (FDR) [24]. We also conducted sample size estimation using the intersected genes among the discovery and validation sets for all possible group combinations. After considering the sample size estimation, the group combinations and the intersected genes were selected for the downstream analysis (Figure 2B).

### 2.3. Transcriptome analysis

Transcriptome analysis was conducted in two stages. In stage 1 (Figure 2B), we used low-level information in each dataset with same experiment and platform to identify a gene set by differential expression analysis. In stage 2 (Figure 2C), high-level information was used across datasets with different experiments and platforms by gene set overlap analysis.

#### 2.3.1. Low-level information analysis to identify differential expression

In stage 1, we only conducted a differential expression analysis for each grouping by utilizing transcriptomic data from the same tissue type, i.e., placenta, and the same platform (Figures 2A and 2B). Before differential expression analysis, we filtered out transcripts that were expressed less than 20^th^ percentile. Transcript expression modelling was conducted. We applied moderated *t*-statistics and multiple testing by Benjamini-Hochberg method. The groups were pairs of the subtypes versus control of microarray data from the same platform. Differentially-expressed transcripts were selected if the FDR was less than 5%. Up- and downregulated transcripts were determined based on positive and negative log_2_ fold change, respectively.

#### 2.3.2. High-level information analysis to identify gene set overlap

In stage 2, since the experiments were conducted with different platforms between the associated factors and the subtypes, to identify association between them we applied association test at high-level information by set operation., i.e., gene set overlap analysis. There were two approaches to overlap a pair of gene sets before applying the Fisher test (Figure 2C): (1) non-regulation-specific overlapping for the region-wise Fisher test; and (2) regulation-specific overlapping for the head-to-head Fisher test. Region-wise Fisher test identified whether an overlap between a pair of gene sets was statistically significant or simply by chance. This test was computed between DEGs of the associated factor with those of the subtype, taking total number of genes of interest into account. Meanwhile, head-to-head Fisher test identified whether co- or counter-expression, indicated by overlaps between a pair of gene sets, were statistically significant or simply by chance (undifferentiated). This test was computed between up- and down regulated DEGs of the associated factor with those of the subtype. If the p-value <0.05, the odds ratio (OR) >1 or and <1 concluded more overlapped DEGs respectively with co-expression and counter expression compared to the opposite regulation. This test determined regulation-specific overlap of interests between the associated factor and the subtype. This approach, however, could not identify the causes of the subtypes; yet, the goal of this study was to gain insights of the possible causes, particularly those related to the pre-pregnancy period and microbial community. For the downstream analysis, we only selected DEGs from the significant non-regulation-specific overlap of interest for each of the subtypes, regardless the significance of the regulation-specific overlap.

### 2.4. Code availability

We used R 4.2.2. To synchronize all the package versions and their dependencies, we used Bioconductor 3.16. All analytical codes are available in https://github.com/herdiantrisufriyana/pec.

## 3. Results

### 3.1. Sample characteristics

From the publicly-accessed microarray datasets collected for the associated factors, the phenotype characteristics were described (Table S3). Leiomyomata and other non-endometrial conditions were found in the endometrium datasets, but this lesion was beyond the tissue of interest. For placenta datasets, the first- and second-trimester samples were taken from pregnant women with gestational ages of respectively 8.43 ± 0 and 16.43 ± 0 weeks on average, which would be compared with those at 41 ± 0 weeks’ gestation on average. Almost all the placenta samples with chorioamnionitis were taken from non-preeclamptic pregnant women without HELLP. Meanwhile, all the placenta samples with HELLP were taken from preeclamptic pregnant women without chorioamnionitis.

The phenotype characteristics were also described for the subtypes (Table S4). No chorioamnionitis was found for placenta samples from pregnant women with PE without or with FGR, regardless of the onset, severity and previous hypertension (i.e., superimposed PE). This situation was the same with the controls of any subtypes, i.e., normotensive pregnancy, and other PIH subtypes, except early-onset gestational hypertension. Pregnant women with HELLP were only found in the severe subtypes of PE, except superimposed, early-onset, severe PE with FGR.

We conducted differential expression analysis within each experiment of the same tissue (Table S5) with 5118 background genes. They were intersected among the microarray datasets with sufficient, gene-wise sample size. Since we did not find a cohort including all the associated factors and the subtypes, we could only identify high-level associations by overlapping the gene sets determined by different cohorts (Figure 2). The role in the placental dysfunction of the PE subtypes was indicated for endometrial maturation but not decidualization (Figure 3), in addition to placentation. Opposite gene regulation was also indicated between the PE subtypes and chorioamnionitis but not between the PE subtypes and HELLP syndrome (Figure 4; Table S6). We also gained insights related to late-onset PE.

**Figure 3.**
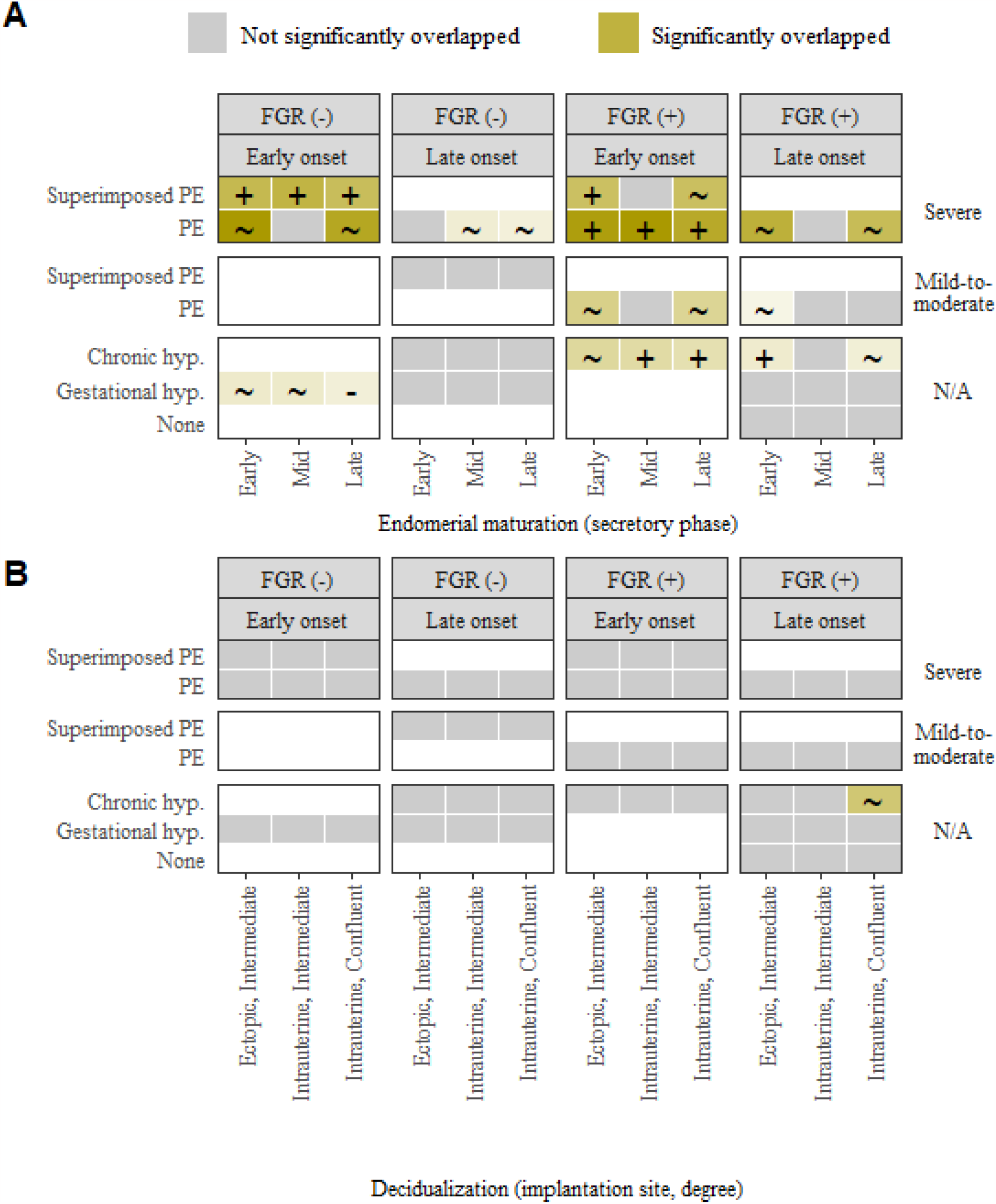
Transcriptome analysis by gene-set overlapping between the subtypes and the associated factors of: (A) Endometrial maturation; and (B) Decidualization. Significances of undifferentiated, co-, and counter-expression are respectively indicated by ∼, +, and -. The color gradation represents the number of overlapped DEGs for undifferentiated expression and the number of either co- or counter-expressed DEGs. All non-grey tiles are significant for the non-regulation-specific overlapping. DEGs, differentially-expressed genes; FGR, fetal growth restriction; hyp.; hypertension; PE, preeclampsia; N/A, not applicable.

**Figure 4.**
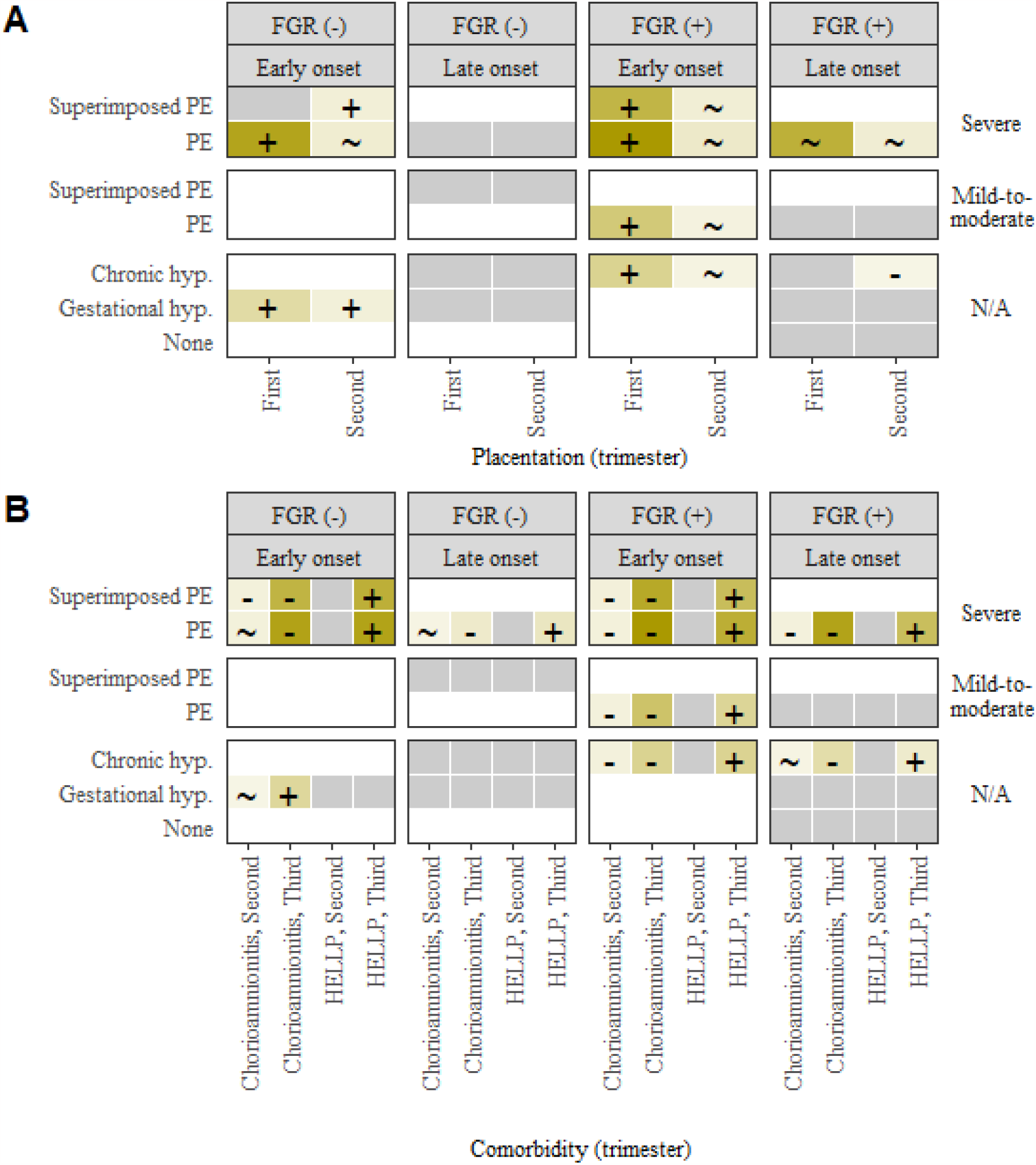
Transcriptome analysis by gene-set overlapping between the subtypes and the associated factors of: (A) Placentation; and (B) Comorbidity. Significances of undifferentiated, co-, and counter-expression are respectively indicated by ∼, +, and -. The color gradation represents the number of overlapped DEGs for undifferentiated expression and the number of either co- or counter-expressed DEGs. All non-grey tiles are significant for the non-regulation-specific overlapping. DEGs, differentially-expressed genes; FGR, fetal growth restriction; HELLP, hemolysis, elevated liver enzymes, and low platelets; hyp.; hypertension; PE, preeclampsia; N/A, not applicable.

### 3.2. Role in placental dysfunction of PE by endometrial maturation but not decidualization

Endometrial maturation, particularly late-secretory phase, showed a potential role in several subtypes. Significant overlaps were found between DEGs of late-secretory endometrium and placenta under any subtypes with: (1) early onset (*p*-values ≤0.008); (2) severe hypertension and proteinuria (*p*-values ≤0.042); or (3) chronic hypertension and/or severe PE with FGR (*p*-values ≤0.042). Among these overlaps, placenta under early-onset, gestational hypertension also indicated significant counter-expression (OR 0.15, 95% confidence interval [CI] 0.04 to 0.44; *p*-value <0.001). Meanwhile, a significant co-expression was identified if the subtypes fulfilled criteria of: (1) early-onset, severe superimposed PE (OR 2.43, 95% CI 1.51 to 3.95; *p*-value <0.001); or (2) early-onset FGR with either severe PE (OR 1.72, 95% CI 1.17 to 2.54; *p*=0.005) or chronic hypertension only (OR 2.85, 95% CI 1.38 to 6.03; *p*-value 0.003) but not both (i.e., superimposed PE; OR 1.15, 95% CI 0.69 to 1.93; *p*-value >0.05). These findings implied endometrial maturation play the putative role by sharing the same up- and down-regulated genes with placenta under those subtypes. Meanwhile, the opposite regulation would lead to gestational hypertension alone without affecting the early onset. In addition, unlike late-secretory endometrium, the overlap patterns were inconclusive between early- and mid-secretory endometrium with the subtypes.

While decidualization is considered a subsequent process of endometrial maturation, our finding did not show its potential role in almost all the subtypes, except late-onset FGR with chronic hypertension. A significant overlap was only found between the DEGs of intrauterine, confluent decidua and placenta under late-onset FGR with chronic hypertension, in which all the decidual DEGs were included in the placental ones (OR ∞, 95% CI 2.09 to ∞; *p*-value 0.025). Neither significant conor counter-expression was identified between both sets of DEGs. Nevertheless, since decidua is a pregnancy version of endometrium, we believe this tissue may connect endometrial maturation and placentation by a process that cannot be identified by gene expression.

Several findings were indeed implying the role of endometrial maturation in placentation. Significant overlaps were also found between DEGs of first- and third-trimester placentas respectively under normotensive pregnancy and any subtypes with chronic hypertension and/or PE with FGR (*p*-values ≤0.035). However, the third-trimester placenta was under neither always early onset nor severe hypertension and proteinuria. This exception differed gene expression of first-trimester placenta from that of late-secretory endometrium in terms of overlaps with placenta of the subtypes. The same overlapping patterns were also applied to second-trimester placenta but there was no regulation-specific overlapping. Meanwhile, significant co-expressions (OR lower bounds ≥1.56) were identified in the aforementioned overlaps of first-trimester placenta. The role of endometrial maturation in placentation might be related to the impact of chronic hypertension and/or PE on fetal growth more than onset and severity.

### 3.3. Competing risk of chorioamnionitis and PE with opposite gene regulation

Furthermore, endometrial maturation implied a putative role in differing PE from chorioamnionitis. Late-secretory endometrium and placenta under chorioamnionitis shared the same subtypes whose placental DEGs overlapped with their respective ones. However, the gene regulations were significantly counter-expressed in majority for chorioamnionitis (*n*=13/18, 72.22%; OR upper bounds ≤0.21). The remaining overlaps were neither conor counter-expressed, except third-trimester placentas under early-onset, gestational hypertension. Its DEGs indicated significant co-expression with those under chorioamnionitis (OR ∞, 95% CI 157.41 to ∞; *p*-value <0.001) but counter-expression with those of late-secretory endometrium (OR 0.15, 95% CI 0.04 to 0.44; *p*-value <0.001). These findings implied that PE and chorioamnionitis might have different endometrial maturation.

### 3.4. Role in HELLP syndrome by endometrial maturation

Similar to placenta under chorioamnionitis, endometrial maturation also implied a putative role in differing PE from HELLP. The similar subtypes were also shared, whose placental DEGs overlapped with those of late-secretory endometrium and placenta under HELLP. However, there was an exception, i.e., early-onset, gestational hypertension. The significant overlaps of HELLP were also only in third-(*p*-values ≤0.001) but not second-trimester placenta. This finding implied that a HELLP syndrome might be a competing risk of PE if it occurs alone in earlier trimester. The role of endometrial maturation in HELLP syndrome might be mediated by PE only. In addition, unlike chorioamnionitis, all the gene regulations were significantly co-expressed in the overlaps between the third-trimester, placental DEGs of the shared subtypes and HELLP syndrome (OR lower bounds ≥12.79). These findings gain insights in differentiating between HELLP that occurs alone and with PE.

### 3.5. Insights related to late-onset, severe PE

Eventually, it is important to point out the difference between any PE subtypes with early and late onset. Most findings up to this point were related to any PE subtypes with early onset. While we identify a significant overlap between the placental DEGs under late-onset, severe PE without FGR and the endometrial ones at mid-(OR 1.7, 95% CI 1.21 to ∞; *p*-value 0.004) and late-secretory phases (OR 1.47, 95% CI 1.06 to ∞; *p*-value 0.026), we did not identify any significant overlaps between these DEGs with the placental DEGs under normotensive pregnancy. Meanwhile, differential expression analysis identified the placental DEGs for: (1) first- and second-trimester versus term placentas; and (2) late-onset, severe PE without FGR versus normotensive pregnancy. These findings implied that placentation did not play any role in late-onset PE without FGR but endometrial maturation did. However, the significant overlaps were identified if this condition was accompanied by FGR (*p*-values ≤0.041); which indicated that placentation might still play a role in FGR under late-onset PE. These findings implied that impaired placentation after impaired endometrial maturation might lead to either an earlier PE or a PE impact on fetal growth.

## 4. Discussion

### 4.1. Interpretation and comparison to previous works

Transcriptome analysis indicated that the role in placental dysfunction of chronic hypertension and/or severe PE with FGR was potentially played by endometrial maturation but not decidualization, particularly gene expression during late-secretory endometrium. The role of endometrial maturation via placentation for these subtypes was considered smaller in affecting their onset and severity. Meanwhile, only the role of endometrial maturation but none of placentation was indicated in late-onset, severe PE without FGR, distinguishing this subtype among others. In addition, endometrial maturation was also indicated to play a role in HELLP syndrome as a subsequent of PE (e.g., during third trimester) but not as its antecedent (e.g., during second trimester). Furthermore, the decision to terminate the pregnancy was likely taken if the preeclamptic pregnant women was accompanied by HELLP, since most HELLP placenta were found in preeclamptic pregnant women without chorioamnionitis. Therefore, endometrial maturation alone in pre-pregnancy period might play a putative role in PE pathogenesis.

For histologic chorioamnionitis, its gene set also overlapped with those of the chronic hypertension and/or severe PE with FGR. It was similar to late-secretory endometrium but the gene regulation was the opposite. While endometrial maturation might also play a role in histologic chorioamnionitis, its gene expression was regulated in the opposite to those subtypes. This finding might be related to the phenotype data, which implied that chorioamnionitis was likely a competing risk of PE. Either PE or chorioamnionitis was probably diagnosed earlier, and in turn, this resulted in early termination before onset of the other condition. Similarly, the competing risk approach have been used for predicting PE with satisfying accuracy [25]. Furthermore, histological changes of placenta in chorioamnionitis were only increased fetal capillaries without villous remodeling as observed in those of PE, but the changes were more acute [26], probably preceding PE. Hence, the gene regulation of endometrial maturation probably determined if a pregnancy would end up with either PE or chorioamnionitis by affecting microbial community in endometrium.

Furthermore, the role of endometrial maturation in pre-pregnancy period was specific to PE among other PIH/PDDs, e.g., early-onset, gestational hypertension. It shared similar characteristics to chorioamnionitis in term of gene regulations which were the opposites to that of late-secretory endometrium. A previous study showed gestational hypertension had the highest risk of acute chorioamnionitis (*n*=29/91, 31.9%; *p*-value <0.001), compared to the other PIH [27]. Nevertheless, what differ early-onset, gestational hypertension from chorioamnionitis is still unclear.

### 4.2. Strength and limitation

By our workflow, an expensive, time-consuming wet lab experiment could be well-prepared not only by literature review but also by a data-driven approach utilizing either at low- or high-level data. This preliminary study also demonstrated how off-the-shelf tools might be variably applied to answer diverse questions in a similar topic. This kind of secondary data analysis across different tissues and timing is inevitable, particularly in pregnancy-related research, because of ethical reasons. Similar situations may also be applied other conditions with long interval time in which a primary data collection is difficult and expensive.

Several limitations are considered in this study. Microarray dataset of gene expression may not help to reveal all parts of the pathogenesis. Nevertheless, we differentiated the several subtypes of PE, and identified the novel, data-driven pathways, using the microarray dataset only. Additional information from the next-generation sequencing data may reveal new perspectives to the proposed pathogenesis of PE in this study, by including non-coding genes. We also could not apply the results directly to develop screening and preventive strategies in clinical settings. This is because we used microarray from tissues by invasive sampling, unlike blood sampling or other methods which are routinely used in clinical settings. Yet, these give a specific direction for the variables and the study design for the next investigation to support the clinical implementation.

## 5. Conclusions

The role in placental dysfunction was potentially played by endometrial maturation, but not decidualization, for any subtypes with early onset, severe hypertension and proteinuria, or chronic hypertension and/or severe PE with FGR. However, no role of placentation was indicated in late-onset, severe PE without FGR. Both phenotype and genotype also implied that histologic chorioamnionitis was likely a competing risk of PE, in which the gene regulation of endometrial maturation might affect surrounding microbial community to determine if a pregnancy ends up with either PE or chorioamnionitis. In addition, our preliminary results showed the feasibility of developing and validating pathogenesis models of PE subtypes which will be the focus of the extension work of this preliminary study. Eventually, this study will help to decide if future studies need prospective, pre-pregnancy and chorioamnionitis data for preeclampsia.

## Acknowledgments

Preprint of an article published in Pacific Symposium on Biocomputing © 2023 World Scientific Publishing Co., Singapore, http://psb.stanford.edu/. We thank Yuan-Chii Lee for assistance with microarray analysis. This study was funded by: (1) the Postdoctoral Accompanies Research Project from the National Science and Technology Council (NSTC) in Taiwan (grant no.: NSTC111-2811-E-038-003-MY2) to Herdiantri Sufriyana; and (2) the Ministry of Science and Technology (MOST) in Taiwan (grant nos.: MOST110-2628-E-038-001 and MOST111-2628-E-038-001-MY2), and the Higher Education Sprout Project from the Ministry of Education (MOE) in Taiwan (grant no.: DP2-111-21121-01-A-05) to Emily Chia-Yu Su.

## Supplementary material

https://github.com/herdiantrisufriyana/pec/blob/master/supplementary_material.pdf.

